# Connectome Principal Component Drives Cross-Dataset Replication and Clinical Prediction in Symptom Lesion Network Mapping

**DOI:** 10.64898/2026.03.04.709716

**Authors:** Sasin Treeratana, Arp-Arpa Kasemsantitham, Setthanan Jarukasemkit, Waragon Phusuwan, Anthipa Chokesuwattanaskul, Sira Sriswasdi, Janine Bijsterbosch, Chaipat Chunharas

**Author notes:** **Corresponding Authors Information** Sasin Treeratana, Chaipat Chunharas.

## Abstract

Lesion network mapping (LNM) describes a group of methods using normative functional connectivity data to map disparate brain lesions and stimulation sites onto common brain networks. Van den Heuvel and colleagues recently showed that these methods lack disease specificity, instead producing maps that converge toward intrinsic properties of the normative connectome dataset. Here, we investigate symptom LNM (sLNM), a recent advancement in the method which attempts to increase the robustness of results by incorporating symptom severity and incorporating replication across multiple datasets and prediction of clinical outcomes. Using clinical datasets of depression and Broca’s aphasia, we show that sLNM maps from unrelated disorders nonetheless converge despite using null models which break the specific lesion–symptom structure in the datasets. Using simulated datasets with a known ground-truth disease network, we show that sLNM results are systematically biased towards the normative connectome’s first principal component (PC1), which drives spurious convergence across unrelated datasets. We further show that the apparent clinical predictive capability of these maps are non-specific: network maps derived from unrelated disorders such as migraine and aphasia predict brain stimulation improvement in depression as well as — or better than — the cohort’s own sLNM map. However, controlling for PC1 reduces spurious convergence across unrelated datasets and improves clinical prediction specificity, supporting the notion that disease-specific signal exists within sLNM but is confounded by the globally present PC1 signal in the normative connectome. These findings offer a practical correction applicable to existing and future sLNM studies.

## 1. Introduction

Lesion network mapping (LNM) is an approach that uses functional connectivity (FC) from a large normative connectome dataset to map focal brain abnormalities in neurological and psychiatric disorders onto common brain networks (Boes et al., 2015; Darby et al., 2019; Fox, 2018). This approach was developed under the premise that focal brain perturbations — such as lesion or brain stimulation — in diverse brain regions may nonetheless produce similar symptoms when they are functionally connected to the same functional network (Darby et al., 2019; Fox, 2018; Siddiqi et al., 2022). Over the past decade, LNM has been applied across numerous brain disorders and behavioral conditions (Joutsa, Corp, et al., 2022; van den Heuvel et al., 2026), including studies proposing new treatment targets for brain disorders (Pines et al., 2025; Siddiqi et al., 2020; Siddiqi, Philip, et al., 2024), as well as prospective clinical trials using LNM-derived transcranial magnetic stimulation (TMS) targets (Cui et al., 2025; J. Taylor et al., 2024).

LNM encompasses various methodological variants (Boes et al., 2015; Burke et al., 2020; Ji et al., 2025; Joutsa, Moussawi, et al., 2022; Peng et al., 2024; Siddiqi et al., 2021; Tetreault et al., 2020), one of the most recent advancements being a symptom-correlation approach, referred to here as **Symptom Lesion Network Mapping (sLNM)** (Ferguson et al., 2024; Goede et al., 2025; Horn et al., 2017, 2022; Li et al., 2021; Milano et al., 2025; Palm et al., 2023; Siddiqi et al., 2020, 2021, 2023, 2025; Siddiqi, Klingbeil, et al., 2024; Siddiqi, Philip, et al., 2024; Sobesky et al., 2022). Whereas conventional LNM identifies brain regions or hubs commonly functionally connected to lesions causing the same clinical symptom, sLNM maps brain regions whose connectivity to lesions predicts the *severity* of symptoms. The same approach can also identify regions whose connectivity to brain stimulation sites predicts clinical improvement (sometimes referred to as ‘*r-maps*’ in the deep brain stimulation (DBS) literature) (Goede et al., 2025; Horn et al., 2017, 2022; Li et al., 2021; Sobesky et al., 2022). Validation of sLNM has relied on replication across multiple datasets and the ability of these maps to predict clinical outcome in independent datasets (Li et al., 2021; Siddiqi et al., 2021, 2026). Furthermore, prospective TMS trials using sLNM-derived stimulation targets specific to anxiety have demonstrated anxiety improvements, supporting the validity and clinical utility of the method (Cui et al., 2025; J. Taylor et al., 2024).

Recently, van den Heuvel and colleagues demonstrated that LNM and its methodological variants, including sLNM, are susceptible to a mathematical artifact in which the output maps converge onto a similar and non-specific spatial pattern regardless of the clinical data used (van den Heuvel et al., 2026). Specifically, conventional LNM was shown to converge toward the degree of the normative connectome — a graph-theoretical metric describing the overall connectedness of each brain region — while sLNM converges toward the first principal component (PC1) of the normative connectome, which reflects a gradient from unimodal to transmodal cortical areas (Margulies et al., 2016; van den Heuvel et al., 2026). These findings raise concerns that LNM maps may not capture disease-specific circuits but are instead shaped by intrinsic properties of the normative connectome data used in the analysis.

The findings by van den Heuvel and colleagues have generated considerable discussion. Several responses have challenged or expanded on van den Heuvel and colleagues’ methodology and conclusions (Edelman et al., 2026; Meng et al., 2026; Petersen et al., 2026; Siddiqi et al., 2026; Zalesky & Cash, 2026). Criticisms include the overlooking of subtle but meaningful differences between LNM maps (Meng et al., 2026; Siddiqi et al., 2026), the use of ecologically invalid lesion datasets (Meng et al., 2026; Siddiqi et al., 2026), and the use of a simplified version of the LNM method (Siddiqi et al., 2026). Some responses have also proposed null models for LNM, including permuting group labels for disease versus control lesions (Petersen et al., 2026), permuting lesion spatial distributions (Zalesky & Cash, 2026), and permuting the FC matrix of the reference connectome itself (Zalesky & Cash, 2026). These null models test whether LNM maps carry information beyond basic properties of the normative connectome, such as hub topology, and the spatial distribution of lesions, by preserving those properties in the null distribution (Petersen et al., 2026; Zalesky & Cash, 2026). However, most of this work has focused on conventional LNM; null models specific to sLNM remain underexplored. Currently, sLNM studies permute symptom labels within disease datasets to break any lesion–symptom relationship, and use replication across independent datasets of the same symptom presentation (or split-half datasets) to test whether sLNM maps across datasets are more spatially similar than expected by chance (Siddiqi et al., 2021; Siddiqi, Philip, et al., 2024). For example, given a dataset of brain lesions causing depression and an independent dataset of brain stimulation targets relieving depression, permutation testing determines whether sLNM maps derived from each dataset are more spatially similar than maps derived from the same datasets with shuffled symptom labels (Siddiqi et al., 2021).

In their study, van den Heuvel and colleagues provided empirical evidence that sLNM maps derived from clinically unrelated datasets (depression and aphasia) are spatially similar, and both closely resemble the connectome PC1 (which also predominantly describes the degree). However, the significance of these similarities was quantified using spatial null methods such as the spin test (Alexander-Bloch et al., 2018) and BrainSMASH (Burt et al., 2020), which attempt to account for spatial smoothness in brain maps, rather than the symptom-label permutation null used in sLNM studies. These spatial null methods have been argued to overestimate the significance of similarity between brain maps (Koussis et al., 2025; Siddiqi et al., 2026). As such, there is an opportunity to leverage symptom-label permutation testing (typically used to establish convergence between independent datasets of the same disease) to determine whether sLNM maps converge between independent datasets of unrelated diseases. Because the current null tests directly test for the *presence* of structured lesion-symptom relationships, not necessarily their *specifics*, it remains theoretically possible for datasets with entirely different underlying lesion-symptom relationships to nonetheless produce convergent sLNM maps. Such unrelated convergence would support the lack of disease-specificity suggested by van den Heuvel and colleagues without the caveats of spatial null models.

In this study, we systematically investigated sLNM. First, we assessed whether sLNM maps systematically converge toward intrinsic features of the normative connectome (PC1), and — because PC1 and degree are highly correlated in real connectome data — we disentangled which of these features drives sLNM outputs. Second, using both the clinical datasets examined by van den Heuvel et al. and semi-simulated datasets with controlled lesion–symptom relationships, we tested whether the current practice of symptom-label permutation can correct for this convergence. Third, we addressed a question with direct clinical implications: if both LNM and sLNM reflect intrinsic connectome features rather than disease-specific circuits, why do maps derived from these methods predict clinical outcomes? We tested the hypothesis that the predictive performance of these maps is itself systematically inflated by connectome PC1. Finally, we demonstrated that controlling for PC1 improves the specificity of both sLNM inference and clinical prediction.

## 2. Methods

Our analyses proceed in several stages. First, we applied the sLNM pipeline to two unrelated clinical datasets — TMS treatment for depression and post-stroke Broca’s aphasia — and tested their convergence using the symptom-label permutation test (**Sections 2.1–2.3**). Second, we used semi-simulated datasets with known ground-truth disease networks to benchmark sLNM’s accuracy in recovering these networks and to characterize whether the pipeline systematically distorts outputs toward connectome features (**Sections 2.4–2.6**). Third, we evaluated the sensitivity and specificity of the symptom-label permutation test under controlled conditions (**Section 2.7**). Finally, we tested whether sLNM maps offer disease-specific clinical predictions in the TMS cohort, or whether maps from unrelated conditions predict outcomes equally well (**Section 2.8**).

### 2.1 Data sources

This study used a healthy control normative connectome dataset to inform sLNM (see **section 2.1.1**) and multiple clinical datasets (see **section 2.1.2**). All data used in this study are publicly available (see data availability statement for further information).

#### 2.1.1 Connectome Dataset

We used the already preprocessed GSP1000 connectome data (Buckner et al., 2020) as the healthy reference cohort for all sLNM analyses. This dataset includes preprocessed resting-state fMRI BOLD time-series from N=1000 subjects (1:1 male:female, age 18–36) in MNI152 template 2-mm isovolumetric space. The voxel-wise seed FC map was calculated by computing seed-to-voxel BOLD temporal correlation (Pearson’s r) for each GSP1000 participant, then averaging correlation values across participants. This was performed using publicly available Python code from a previous LNM study (https://github.com/nimlab/NHB_Taylor2023; J. J. Taylor et al., 2023). Seeds were defined from the clinical datasets as described below.

#### 2.1.2 Clinical Datasets

We used two publicly available clinical datasets. The first dataset (Boston TMS cohort) consists of N=25 subjects with depression who underwent a TMS clinical trial (Weigand et al., 2018). The TMS target sites were recorded as MNI coordinates and the outcome of interest was percent reduction in Beck Depression Inventory (BDI) scores. 4-mm radius spheres were generated at each reported TMS site and used as seeds for subsequent analyses.

The second dataset used post-stroke Broca’s aphasia subjects from the Aphasia Recovery Cohort (ARC), an open-source chronic stroke data repository (Gibson et al., 2024). Segmented and preprocessed lesion seed files as well as de-identified subject clinical data were downloaded from the ARC GitHub repository (https://github.com/neurolabusc/AphasiaRecoveryCohortDemo). Of the 86 subjects with Broca’s aphasia, one was excluded due to a missing preprocessed lesion mask, resulting in a final sample of N=85. Here the outcome of interest was the post-stroke Western Aphasia Battery Aphasia Quotient (WAB-AQ).

### 2.2 Symptom Lesion Network Mapping

The sLNM pipeline was performed as described in (Siddiqi et al., 2020). For a given dataset, each subject’s seed (lesion or TMS-site) was mapped to a voxel-wise mean FC map using the GSP1000 dataset. Seed FC values were correlated (Pearson’s r) with relevant clinical outcomes at each voxel across subjects, producing an sLNM map for that dataset. Therefore, each voxel in the output map encodes the correlation value between FC strength to lesion or TMS-site and symptom severity. Applying this pipeline to Dataset 1 (Boston TMS cohort) and Dataset 2 (ARC-Broca’s aphasia) produced the Depression-TMS network, and Broca’s aphasia network, respectively (**Fig. 1A**).

**Figure 1.**
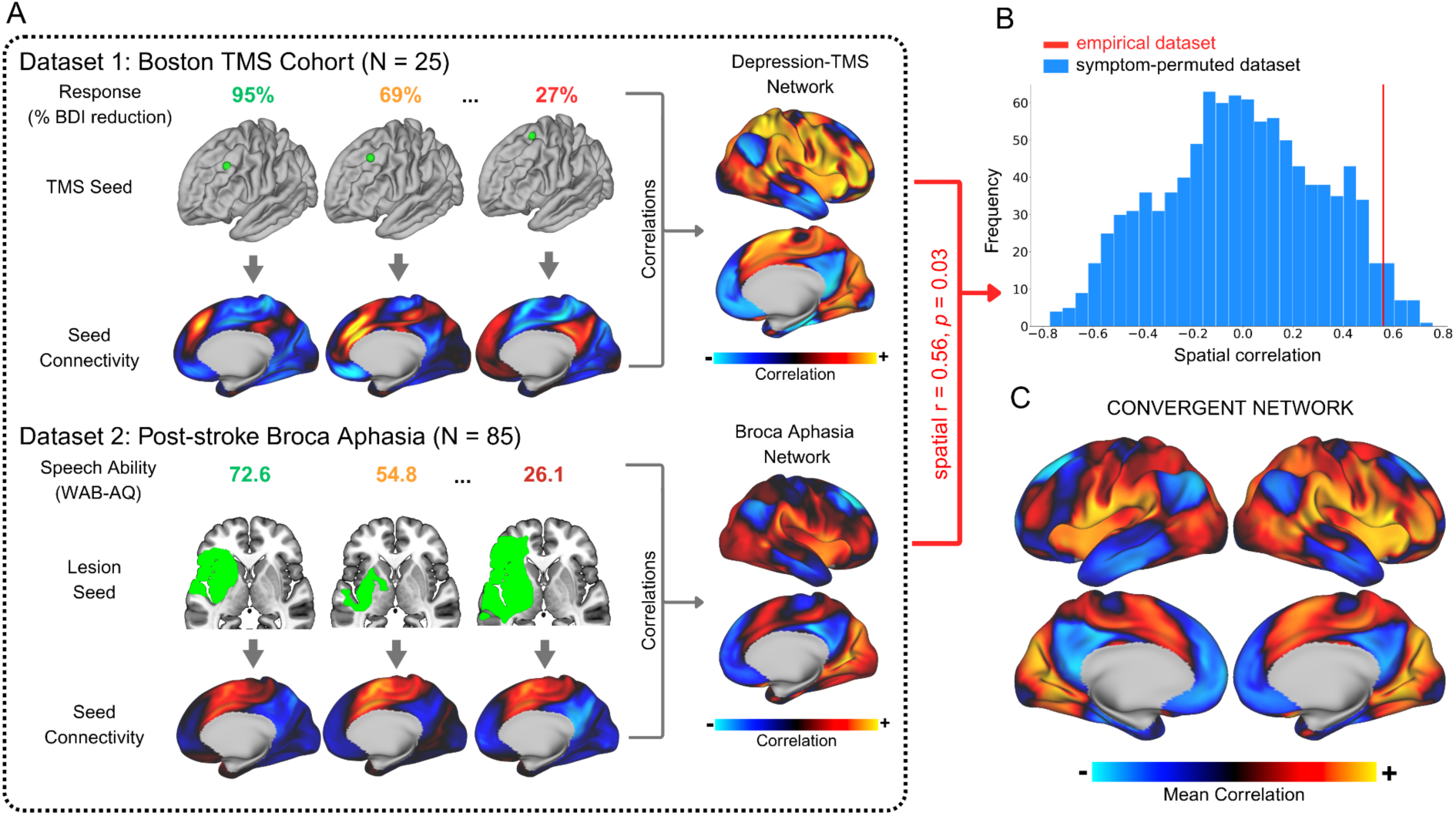
**A)** Schematic of sLNM analysis. Seed regions were mapped onto mean voxel-wise FC maps using the GSP1000 connectome. Seed FC values were then correlated with clinical outcomes at each voxel to produce sLNM maps. This was applied to the Boston TMS cohort (N=25) and the ARC Broca’s aphasia cohort (N=85), yielding the Depression-TMS Network and Broca’s Aphasia Network, respectively. **B)** Symptom-label permutation test. The spatial correlation between the two networks was r = 0.56. A null distribution was generated by shuffling clinical outcome labels within each dataset and recomputing the spatial correlation across 1000 permutations. The observed convergence was significant (p = 0.03). **C)** Convergent network. The two sLNM maps were averaged into a single composite map, proposed to explain both depression response to TMS and Broca aphasia. Voxel-wise statistical maps were visualized in fsLR-32k surface space.

### 2.3 Symptom-label Permutation Testing

To assess whether two datasets converged onto a common brain network, we first computed sLNM maps for both datasets, then, the spatial correlation (Pearson’s r) between the two datasets’ sLNM maps was computed to determine similarity. Then, within each dataset, symptoms were shuffled so that each patient’s lesion was paired with another patient’s symptom score, and correlations between permuted sLNM maps from the two datasets were recalculated to generate a null distribution (1000 permutations). The p-value was defined as the proportion of permuted spatial correlation equal to or exceeding the observed value, similar to previous studies (**Fig. 1B**) (Joutsa, Moussawi, et al., 2022; Milano et al., 2025; Palm et al., 2023; Siddiqi et al., 2020, 2021, 2023; Siddiqi, Klingbeil, et al., 2024; Siddiqi, Philip, et al., 2024).

### 2.4 Simulating Lesion-Symptom Datasets

To generate simulated datasets with paired lesion and symptom score data, we designed a model inspired by previous sLNM studies (Ferguson et al., 2024; Goede et al., 2025; Horn et al., 2017; Li et al., 2021; Siddiqi et al., 2021, 2023, 2025; Sobesky et al., 2022). In those studies, each subjects’ clinical outcome was predicted by the spatial similarity between their lesion or stimulation seed connectivity profile and a pre-established disease network (for a clear example, see *Figure 1* in Goede et al., 2025). Our semi-simulated model mirrors this structure (**Fig. 2A**).

**Figure 2.**
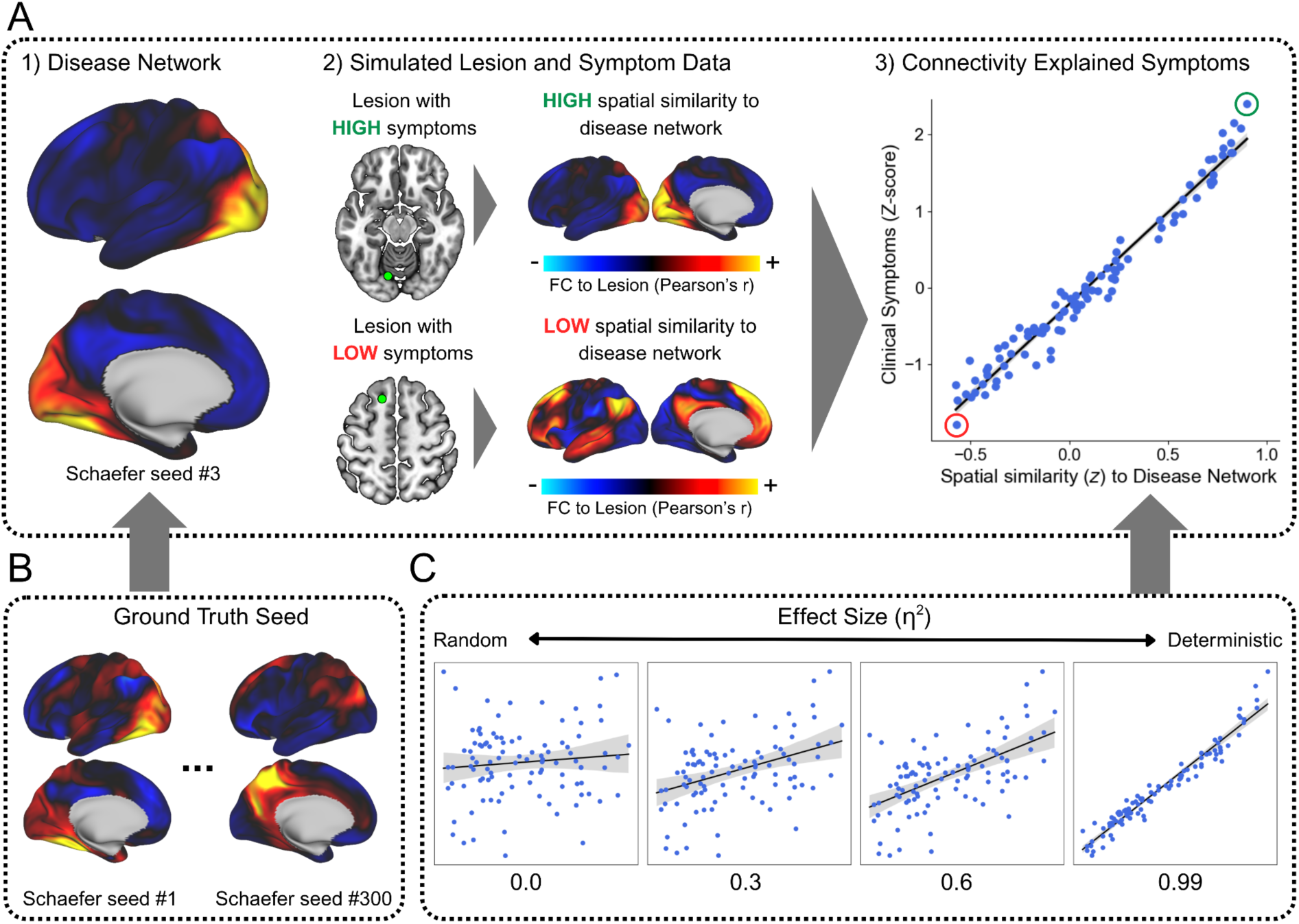
**A)** Simulation framework. Each lesion was defined as a 4-mm radius sphere placed randomly within the brain. The ground-truth disease network was defined as the FC map of a randomly selected cortical seed from the 300-region Yeo-Schaefer atlas (seed #3 in the left visual network is shown as an example). To model symptoms, each subject’s lesion FC map was computed from the mean connectome, and its spatial similarity (Fisher-z transformed Pearson’s r) to the ground-truth network determined the symptom score. **B)** Examples of other Schaefer seed FC maps that could be used as the ground-truth network instead of the one in A. **C)** Effect size settings determining the lesion–symptom relationship: random (η² = 0.0), realistic (0.3), strong (0.6), and deterministic (0.99). For each plot, the x-axis shows the z-transformed spatial correlation between each lesion FC map and the ground truth; the y-axis shows the symptom score (z-scored). The shaded area represents the 95% confidence interval. All voxel-wise statistical maps were reconstructed and visualized on the fsLR-32k surface.

The steps to generate a synthetic dataset were as follows: first, a set of lesions was generated (N=100); details on lesion generation can be found in the next section. Then, to pair each lesion with a symptom score, we assigned a ground-truth network. To ensure biological plausible topography, ground-truth networks were defined as FC maps of a randomly selected cortical parcel from the Yeo–Schaefer 300-parcellation atlas (Schaefer et al., 2018), yielding 300 unique possible ground-truth networks (**Fig. 2B**). Lastly, we calculated the expected symptom-score for each lesion. For each lesion, spatial similarity to the ground-truth network (*R*) was defined as the Fisher-z transformed spatial Pearson’s r between the lesion’s FC map and the ground-truth network. Symptoms (*S*) were then modeled as a linear function of *R* plus noise (*ε*) and subsequently z-scored:

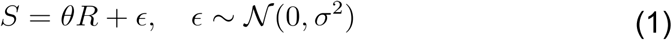

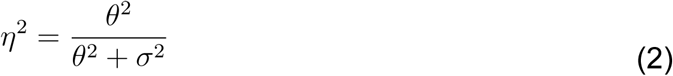

The effect size parameter (*η^2^*) controlled the proportion of symptom variance explained by the ground-truth network, tested at four levels, representing lesion-symptom relationships that are random (0.0), weak (0.3), strong (0.6), and deterministic (0.99) (**Fig. 2C**). **Figure 2A** demonstrates a simulated dataset with a visual region (Schaefer seed #3) as seed for the ground truth disease network, with effect size of 0.99.

This simulation approach differs fundamentally from previous LNM simulations, which manipulate lesion characteristics such as size, number, and spatial distribution to test how these factors affect pipeline outputs. In those studies (van den Heuvel et al., 2026; Zalesky & Cash, 2026), the ground truth defines how lesions are distributed, for example, lesions are placed preferentially within the default mode network (DMN) versus outside it. Such approaches are appropriate for conventional LNM, where the output is determined by the spatial distribution of lesions. In contrast, sLNM incorporates symptom measures — the output map encodes correlations between lesion connectivity and symptom severity, not simply lesion distribution. Consequently, given a fixed set of lesions, sLNM produces different maps depending on how symptoms are paired with those lesions. Our simulations test this specific property: rather than varying lesion distributions, we use uniformly distributed lesions and instead define the ground truth at the level of the symptom-generating process — specifying which disease network drives the relationship between lesion and clinical outcome. This directly evaluates whether sLNM can recover the correct disease network influencing symptom data, without confounding the results with assumptions about lesion geometry.

### 2.5 Assessing the Accuracy of sLNM in Recovering Ground Truth

Because each simulated dataset has a known ground-truth network used to generate the lesion-symptom relationship, we can directly benchmark the accuracy of sLNM in recovering that ground truth. We first generated 100 random lesions as 4-mm radius spheres uniformly sampled across all brain voxels. Using the same lesion set but varying the ground-truth network and the effect size of the lesion-symptom relationship, we generated different synthetic clinical datasets that share identical lesion sets but differ in their symptom profiles. Specifically, we paired the fixed lesion set (N=100) with each of 300 Yeo-Schaefer region seed maps as ground-truth networks at four effect sizes outlined previously (0.0, 0.3, 0.6, 0.99), yielding 1200 datasets in total. The goal of sLNM — which takes lesion and symptom scores as input — is to accurately recover the ground-truth network which explains the lesion-symptom relationship.

We evaluated recovery accuracy in two ways. First, we computed the spatial similarity (Pearson’s r) between each ground-truth network and its corresponding sLNM map across effect sizes. Higher effect sizes should yield stronger spatial correlations, while at zero effect size — where symptoms are independent of lesions — the ground-truth network and the sLNM map should be uncorrelated. Second, because spatial correlation is a single measure of global similarity that may mask subtle differences between brain maps (Zalesky & Cash, 2026), we examined the *alignment* of both the ground-truth network and the sLNM map with the first three principal components (PCs) of the GSP1000 connectome, quantified as the absolute Fisher-Z transformed spatial correlation with each PC. This analysis was motivated by empirical results from van den Heuvel and colleagues, suggesting that sLNM output converges toward connectome PC1. Here, principal component analysis (PCA) was performed on the GSP1000 dataset using *IncrementalPCA* (scikit-learn; Pedregosa et al., 2011), which processes data in sequential batches to iteratively update the estimated PCs (one subject per iteration), avoiding the need to compute the full voxel-wise covariance matrix (approximately 292k × 292k dimension), a step that is computationally expensive at this scale. This yields a voxel-wise whole-brain PC map (**Fig. 3B**), rather than the cortical-surface-only or parcellation-based PCs typically derived from group-averaged connectome matrices (Bolt et al., 2022; van den Heuvel et al., 2026).

**Figure 3.**
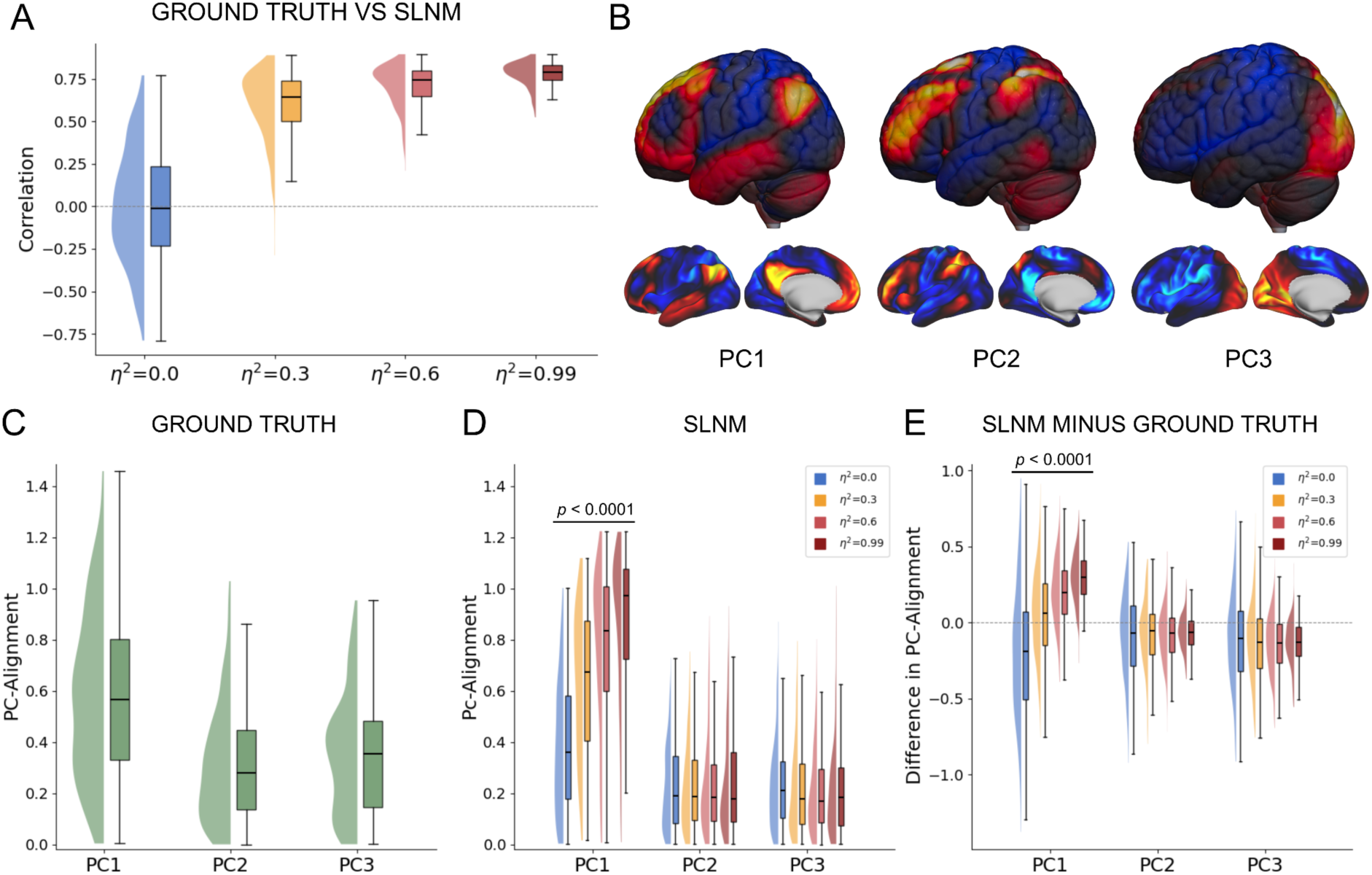
**A)** sLNM correlation to ground truth. Across four effect sizes, sLNM maps showed increasing spatial correlation with the ground-truth network. **B)** Voxel-wise and surface visualization of the first three spatial PCs of the GSP1000 connectome. **C)** Alignment (|z|) of each ground-truth network (N = 300) with PC1–3 as a baseline. **D)** Alignment (|z|) of each sLNM map with PC1–3 across all four effect sizes, showing a progressive increase in PC1 alignment at higher effect sizes (Jonckheere-Terpstra p < 10⁻⁴). **E)** Pairwise difference in PC-alignment (PC1–3) between sLNM and ground truth. PC1 alignment increased with effect size (Jonckheere-Terpstra p < 10⁻⁴), indicating that sLNM systematically shifts map geometry toward PC1 beyond what is present in the ground truth.

To test whether PC alignment increased monotonically with lesion–symptom effect size (e.g., stronger PC1 alignment at higher η²), we applied the Jonckheere-Terpstra test to the PC alignment values of sLNM maps across the four effect-size levels. Significance was assessed by comparing the observed test statistic against a null distribution generated by 10,000 random permutations of the effect-size labels across simulation runs.

Finally, to quantify how sLNM systematically transforms the ground-truth signal in the connectome’s latent space, we then computed, for each PC (1–3), the difference between the sLNM and ground-truth PC alignment values. Positive values indicate that sLNM overrepresents that component relative to the ground truth, and negative values indicate underrepresentation. Monotonic trends in these differences across effect sizes were assessed using the Jonckheere-Terpstra test described above.

### 2.6 Dissociating PC1 and Degree in sLNM Outputs

Continuing from the previous section, because the GSP1000 degree map and PC1 are spatially similar (empirically spatial |r| = 0.54 for whole-brain voxel-wise map, |r| = 0.84 for cortical surface-only map), both highlighting similar regions such as default-mode network nodes, it is challenging to determine which factor drives sLNM results. As such, we performed simulation analyses where we explicitly dissociate the PC1 map from the degree map.

First, we downsampled all analyses from voxel space (approximately 292k voxels) to 1000 Yeo-Schaefer parcellation atlas to enable connectome matrix randomization simulations. To generate the group connectome matrix (C), we computed the parcel-parcel FC values and averaged across all GSP1000 subjects. Then, we decomposed C into eigenvectors and eigenvalues, corresponding to principal components and their explained variance, respectively. Meanwhile, the degree map was calculated as the row sum of C. By swapping the first and k-th eigenvalues in the eigenvalue matrix, we attributed the most variance to the k-th principal component, producing a version of the original connectome with a different PC1 (C_k_). We then calculated the row sum of C_k_ corresponding to its degree. This randomization served two purposes: (1) to dissociate degree and PC1 of Ck, allowing us to see which structure sLNM converges toward, and (2) to determine whether the convergence is mathematical or driven by a biologically valid topographic pattern in PC1 (**Fig 4**).

**Figure 4.**
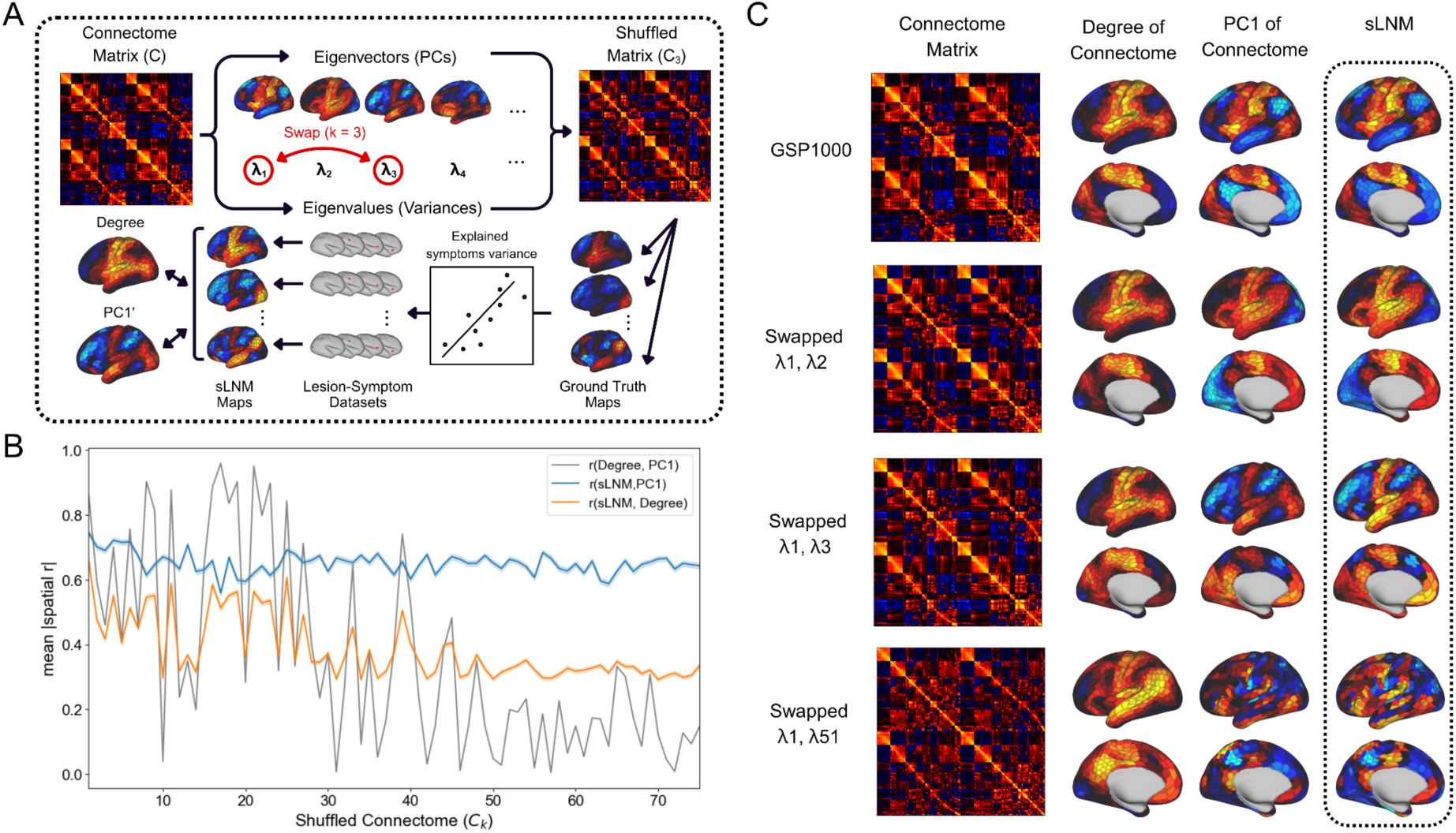
**A)** Connectome randomization procedure. The group-averaged GSP1000 connectome matrix (C; 1000 × 1000 Schaefer parcellation) was eigendecomposed into eigenvectors (PCs) and eigenvalues. To test whether sLNM tracks PC1 specifically, the k-th eigenvalue was swapped with the 1st, assigning the greatest variance to the k-th PC and making it the new dominant component. For each shuffled matrix (C_k_), 1000 datasets were generated from 1000 ground-truth networks, and the resulting sLNM maps were compared with both the degree map and PC1 of C′. **B)** Mean absolute spatial correlation of sLNM with PC1 and degree across 75 shuffled connectomes (N=1000 sLNM maps per connectome). sLNM consistently tracked PC1, whereas its similarity to degree varied widely and covaried with the degree-PC1 correlation, indicating that PC1 — not degree — is the primary driver of sLNM output. Shaded area represents standard error of the mean. **C)** Visualization of the connectome matrix, its degree map, its PC1, and the dominant sLNM pattern (PC1 weights from PCA of 1000 sLNM maps) for selected shuffled connectomes.

For each randomized matrix C_k_, we repeated the analysis described previously (**section 2.5**) by simulating a dataset with a fixed set of 100 lesions. Here, each lesion was a single random cortical (Schaefer-1000) region, with their FC map equal to the corresponding row in C_k_. Symptoms associated with the lesion were modeled using various ground-truth networks with effect size η² = 0.99; this effect size was chosen to ensure that any systematic bias in sLNM outputs reflects the mathematical structure of the pipeline rather than noise. We systematically used each of the 1000 Schaefer parcels’ FC maps as a ground-truth network, yielding 1000 sLNM maps (one per ground truth network) which were compared against the degree and PC1 of C_k_ using absolute spatial correlation (**Fig 4A**). Analysis was performed up to k=75 connectomes for a total of 75,000 sLNM maps.

### 2.7 Sensitivity and Specificity of the Symptom-label Permutation Test

The symptom-label permutation test (**section 2.3**) is used to infer whether two or more datasets share the same underlying disease network explaining lesion–symptom relationships. Here, we evaluate the sensitivity and specificity of this procedure by performing 4000 simulated sLNM studies (1000 studies at each of four effect size levels, as described in **section 2.6**). Each simulated study consisted of two datasets with 100 lesions each, sampled with replacement from a pool of 500 spherical lesions (4-mm radius) distributed uniformly across the brain. Critically, we randomly selected two different ground truth networks for each dataset by choosing two distinct Yeo–Schaefer atlas seed maps. As a result, the two datasets may share highly correlated ground truth networks (seed maps belonging to the same Yeo network), anticorrelated networks (seed maps from anticorrelated Yeo networks, such as the default mode and sensorimotor networks), or uncorrelated networks (such as sensorimotor and visual network seeds). The effect size determined how strongly these ground truth networks influenced the lesion–symptom relationship. For each dataset, we computed the sLNM map and tested the spatial similarity between the two maps using the symptom-label permutation test (significance threshold at p < 0.05). We expected simulated studies whose datasets share related ground truth networks (high spatial r) to produce high significance rates, while those with unrelated ground truth networks (near zero spatial r) should yield significance rates near the nominal 5% level. **Figure 5A** illustrates the schematic: two datasets were generated from unrelated ground truth networks (r = 0.01) at effect size η² = 0.99, and the resulting sLNM maps were tested for convergence to assess whether the permutation test correctly rejects similarity when the underlying disease networks are known to differ.

**Figure 5.**
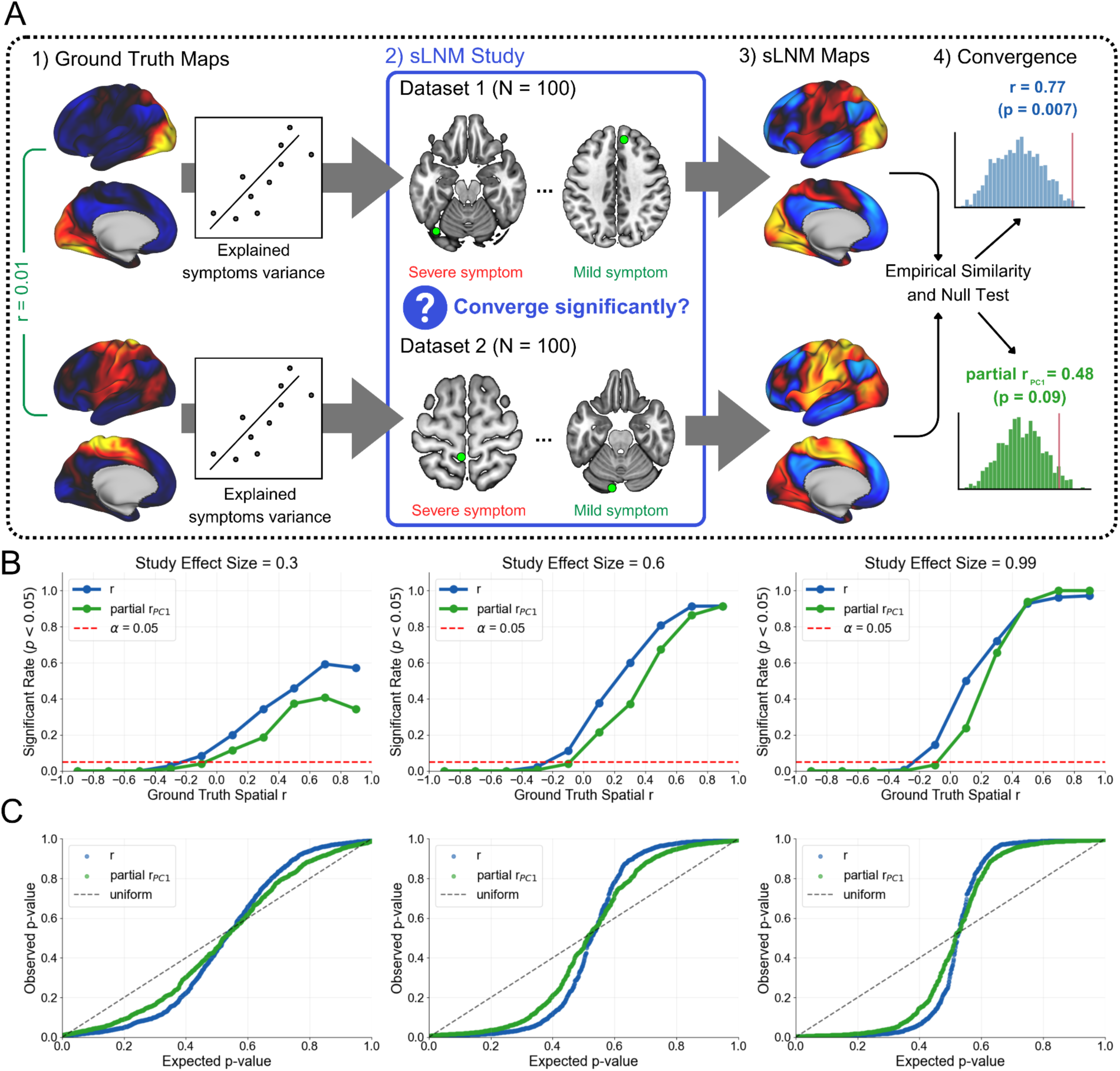
**A)** Simulated sLNM convergence study. For each simulated study, two datasets (N = 100 each) were generated from different ground-truth disease networks, representing two distinct clinical cohorts tested for network convergence. sLNM was performed on each dataset separately, and the spatial similarity of the resulting maps was assessed using the standard symptom-label permutation test. An alternative test using partial correlation controlling for PC1 was also evaluated. The example shown uses two dissimilar ground-truths (r = 0.01) at η² = 0.99: the standard test yielded spurious convergence (r = 0.77, p = 0.007; 1000 permutations), whereas the PC1-controlled test did not (partial r = 0.48, p = 0.09). **B)** Significance rate as a function of ground-truth similarity. Significance rate increased with ground-truth similarity across all effect sizes. At low or zero ground-truth similarity, the standard test produced spurious significance rates exceeding the 5% nominal threshold, with inflation more pronounced at higher effect sizes. The PC1-controlled test reduced these false positives while preserving sensitivity for truly similar networks at higher effect sizes. **C)** Quantile-quantile (QQ) plot of p-values. P-values across all simulated studies followed a bimodal distribution concentrated near zero and one, most prominently at higher effect sizes.

Finally, we tested whether controlling for connectome principal components affects the outcome of the permutation test. Instead of using the spatial correlation between sLNM maps as the test statistic, we used the partial spatial correlation controlling for connectome PC1, computed as:

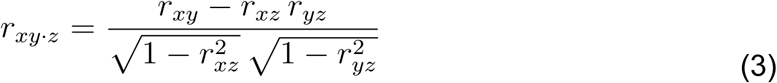

where *x* and *y* are the two sLNM maps whose similarity is being tested and *z* is the PC1 map. This formulation is mathematically identical to regressing out PC1 from both maps and computing the Pearson correlation between their residuals. The same partial correlation coefficient was computed for each permutation to generate the null distribution. If PC1 accounts for spurious convergence between unrelated datasets, using this partial correlation should reduce false positive rates for unrelated ground-truth networks while leaving convergence rates for genuinely related networks intact. Lastly, as controls, we performed the same partial correlation controlling for PC2 or PC3. Because these are not expected to drive the bias, controlling for them should have little effect on the results.

### 2.8 Predicting Clinical Outcomes

To test the clinical utility and specificity of sLNM maps, we performed leave-one-subject-out analyses to predict TMS response in the Boston TMS cohort. For each left-out subject, an sLNM map was generated from all remaining subjects. The Fisher z-transformed spatial correlation between the left-out subject’s TMS-seeded FC map and the leave-one-subject-out sLNM map was then used to predict their TMS response, as measured by percent reduction in BDI (**Fig. 6A**), similar to previous studies (Ferguson et al., 2024; Goede et al., 2025; Horn et al., 2017; Li et al., 2021; Siddiqi et al., 2021, 2023, 2025; Sobesky et al., 2022).

**Figure 6.**
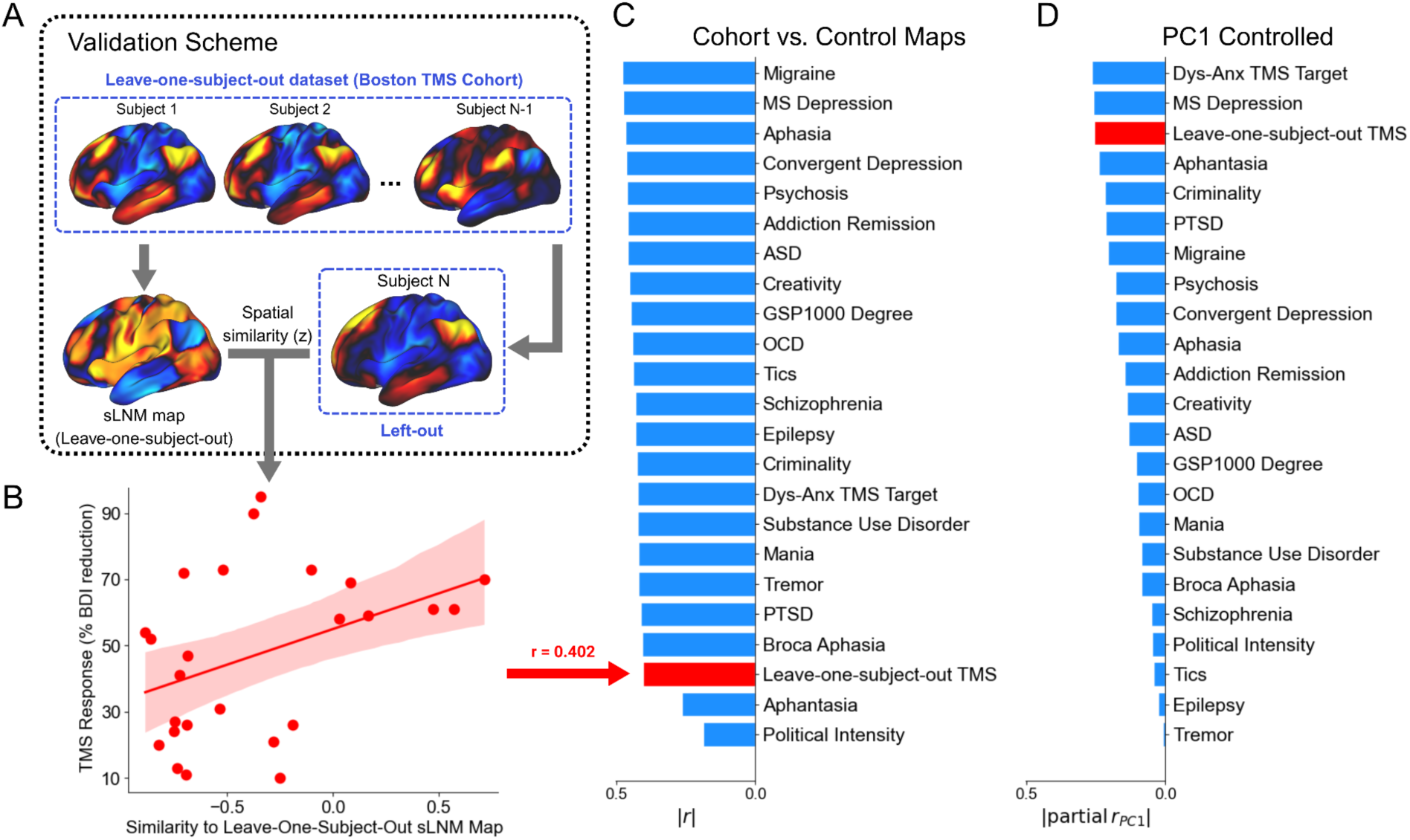
**A)** Clinical validation scheme. Leave-one-subject-out cross-validation was applied to the Boston TMS cohort. For each left-out subject, the sLNM map was recomputed from the remaining subjects. **B)** The spatial correlation between this map and the left-out subject’s TMS-seeded FC map was used to predict TMS response (|r| = 0.402). The shaded area represents the 95% confidence interval. **C)** Non-specificity of sLNM prediction. Control maps derived from unrelated clinical conditions matched or exceeded the prediction of the cohort’s own sLNM map for TMS response, indicating that predictive performance is not disease-specific. **D)** Effect of PC1 correction on prediction. Controlling for spatial similarity to PC1 during prediction reduced the predictive performance of the cohort’s own sLNM map (|r| = 0.256), though it remained among the strongest predictors. Many unrelated maps that had previously outperformed the cohort’s map dropped to near-zero prediction after correction, suggesting that the PC1 signal shared across sLNM and LNM maps is responsible for both inflated predictive performance and apparent cross-disease prediction. ASD, autism spectrum disorder; Dys-Anx, dysphoric-anxiosomatic; MS, multiple sclerosis; OCD, obsessive-compulsive disorder; PTSD, post-traumatic stress disorder

To test the specificity of the method, we additionally performed the same analysis based on spatial correlation between each subject’s TMS-seeded FC map and maps from a range of disorders, sourced directly from previous LNM and sLNM studies, as well as maps available in the GitHub repository of van den Heuvel and colleagues (https://github.com/dutchconnectomelab/lesionnetworkmapping/). This included maps from previous studies or conditions, including migraine (Burke et al., 2020), depression secondary to multiple sclerosis (MS-Depression; Siddiqi et al., 2023), lesions and brain stimulations associated with depression (convergent depression; Siddiqi et al., 2021), aphasia (Zarifkar et al., 2023), psychosis (Pines et al., 2025), addiction remission (Joutsa, Moussawi, et al., 2022), autism spectrum disorder (ASD; Segal et al., 2023), creativity (Kutsche et al., 2025), obsessive-compulsive disorder (OCD; Segal et al., 2023), tics (Ganos et al., 2022), schizophrenia (Segal et al., 2023), epilepsy (Ji et al., 2025), criminality (Darby et al., 2018), dysphoric-anxiosomatic TMS target (Siddiqi et al., 2020), substance use disorder (Stubbs et al., 2023), mania (Cotovio et al., 2020), tremor (Joutsa et al., 2018), post-traumatic stress disorder (PTSD; Siddiqi, Philip, et al., 2024), Broca’s aphasia (computed from the ARC dataset), aphantasia (Kutsche et al., 2026), and political intensity (Siddiqi et al., 2025). Additionally, we included the degree map (van den Heuvel et al., 2026), which was derived entirely from the healthy control GSP1000 connectome with no patient information. For additional descriptions and visualizations of these maps, see **Supplementary Table S1-3** and **Fig. S3**, respectively. Because these maps were all derived from independent data, leave-one-subject-out cross-validation was not required; instead, the spatial correlation between each subject’s TMS-seeded FC map and each control map was used directly to predict their TMS response. If sLNM produces disease- or dataset-specific networks, then maps from unrelated disorders and maps derived solely from the healthy connectome should predict clinical outcomes poorly.

Alternatively, if these control maps predict TMS response as well as the cohort leave-one-subject-out sLNM map, their predictive value cannot reflect disease-specific information. We hypothesized that this non-specific predictive value arises from each map’s similarity to PC1. To test this, we computed a prediction value controlling for PC1 using equation (3), where *x* is the TMS response, *y* is the Fisher-z transformed spatial correlation between each subject’s TMS-seeded FC map and the predictor map, and *z* is the Fisher-z transformed spatial correlation between each subject’s TMS-seeded FC map and the PC1 map. This partial correlation tests whether the predictor map explains clinical outcomes independent of PC1. If a map’s predictive value is driven by PC1 rather than disease-specific spatial information, controlling for PC1 should substantially reduce the prediction, whereas controlling for PC2 or PC3 should have little effect.

## 3 Results

### 3.1 Unrelated Disorders Converged onto a Common Brain Network

Despite being unrelated disorders with different etiologies and clinical presentations, the ‘Depression-TMS’ and ‘Broca’s Aphasia’ networks showed relatively high spatial similarity (r=0.56), which was significant after symptom-label permutation (p=0.03; 1000 permutations) (**Fig. 1B**). This confirms that the networks predicting symptoms in each dataset were more spatially similar than expected by chance.

### 3.2 Connectome Spatial Principal Component 1 Artifact in sLNM Maps

To assess whether sLNM faithfully recovers a known ground-truth network, we applied the method to simulated datasets where the ground-truth disease network was specified by design. Using the same fixed lesion set but varying the ground-truth network and effect sizes which determined lesion-symptom relationships, we found that sLNM produces maps with higher spatial similarity to the ground truth network at higher effect sizes (mean spatial r ± SD = 0.00 ± 0.33 at η² = 0.0, 0.60 ± 0.19 at η² = 0.3, 0.71 ± 0.12 at η² = 0.6, and 0.78 ± 0.07 at η² = 0.99) (**Fig. 3A**).

While sLNM recovered the global structure of the ground-truth network (measured by spatial correlation), the resulting maps were disproportionately aligned with connectome PC1, and this alignment increased monotonically with effect size (Jonckheere-Terpstra p < 10⁻⁴: mean PC-alignment ± SD = 0.37 ± 0.24 at η² = 0.0, 0.66 ± 0.30 at η² = 0.3, 0.78 ± 0.29 at η² = 0.6, and 0.86 ± 0.28 at η² = 0.99).

No such trend was observed for PC2 (Jonckheere-Terpstra p = 0.67) or PC3 (Jonckheere-Terpstra p = 0.65).

Subtracting each ground-truth network’s PC-alignment from that of its corresponding sLNM map confirmed that the growing PC1-alignment was attributable to sLNM itself, not to PC1 structure already present in the ground-truth network. At η² = 0.0, which simulates datasets with no true lesion–symptom relationship (equivalent to shuffled symptom labels), sLNM maps showed slight underrepresentation of PC1 relative to ground truth (mean ΔPC-alignment ± SD = −0.20 ± 0.41). However, when a meaningful lesion–symptom relationship was present, PC1 overrepresentation emerged and scaled with effect size (Jonckheere-Terpstra p < 10⁻⁴: mean ΔPC-alignment ± SD = +0.08 ± 0.26 at η² = 0.3, +0.20 ± 0.19 at η² = 0.6, and +0.28 ± 0.17 at η² = 0.99). In contrast, PC2 and PC3 showed slight decreases in alignment that remained stable across effect sizes (PC2: Jonckheere-Terpstra p = 0.55; PC3: Jonckheere-Terpstra p = 0.63).

In other words, the stronger the true lesion–symptom relationship, the more the sLNM map resembled the ground truth globally (as measured by spatial correlation) but with amplified PC1 features at the expense of other higher-order PCs. To visualize this effect directly, Supplementary **Fig. S1** shows, for each PC, the raw correlation of every ground-truth network plotted against the raw correlation of its corresponding sLNM map. The plot reveals a clear, effect-size-dependent amplification of PC1, while PC2 and PC3 remain close to the ground truth.

### 3.3 Principal Component — not Degree — Accounts for sLNM Output

To disentangle the contributions of PC1 and degree to sLNM outputs, we generated 75 randomized connectomes in which the topography of PC1 and the degree map were dissociated (**Fig. 4A**). For each randomized connectome, sLNM maps consistently showed higher spatial correlation with PC1 than with degree (mean |r| = 0.65 ± 0.03 vs. 0.39 ± 0.10), and sLNM-degree similarity tracked the degree-PC1 correlation of the randomized connectome rather than reflecting an independent relationship (**Fig. 4B**). These results confirm that PC1 drives sLNM outputs rather than degree (**Fig. 4C**), distinguishing sLNM from conventional LNM, which has been shown to converge toward the degree of the connectome (van den Heuvel et al., 2026), while also confirming that the convergence is mathematical rather than biological.

### 3.4 Controlling for PC1 Improves Specificity of Symptom-Label Permutation Test

To test whether the PC1 bias inflates significance rates, we ran simulated sLNM studies, each containing two distinct datasets, and quantified the significance between the sLNM maps using symptom-label permutation testing (**Fig. 5A**). First, we validated the test on null data using 1000 simulated sLNM studies where lesion–symptom relationships were random (η² = 0.0). Here, 3.9% of studies produced significantly similar sLNM maps (at p < 0.05), with a uniform p-value distribution (Kolmogorov-Smirnov test, D = 0.04, p = 0.16), indicating a proper null test for a completely random dataset.

As expected, simulated studies whose two datasets shared highly similar ground truth networks (spatial r = 0.8–1.0) showed high significance rates, reaching 57.1% at an effect size of 0.3, 91.4% at 0.6, and 97.1% at 0.99, confirming that the test is sensitive to true convergence (**Fig. 5B**). Yet when the two datasets had low ground-truth similarity (spatial r = 0.0–0.2), significance rates were substantially inflated above the nominal 5%: 20.0% at η² = 0.3, 37.7% at η² = 0.6, and 50.0% at η² = 0.99. Repeating the analysis controlling for PC1 reduced significance rates at low ground-truth similarity: from 20.0% to 11.5% at η² = 0.3, from 37.7% to 21.5% at η² = 0.6, and from 50.0% to 23.8% at η² = 0.99. Meanwhile, significance rates for studies with genuinely related ground-truths remained largely intact; at high ground-truth similarity (spatial r = 0.8–1.0), both full and partial tests reached >90% significance rates at strong effect size studies (η² ≥ 0.6), though sensitivity was reduced for weak effect size studies (η² = 0.3).

The p-value distribution was bimodal, with most values concentrated near zero or one (**Fig. 5C**). Because sLNM maps project strongly onto either the positive or negative pole of PC1 (**Fig. S1**), two maps are either strongly positively correlated (when their PC1 projections share the same sign) or strongly negatively correlated (when their signs are opposite). Both cases produce extreme test statistics, pushing p-values toward either end of the distribution. Controlling for PC2 or PC3 affected neither the significance rates nor the p-value distribution (Supplementary **Fig. S2**).

Together, these results demonstrate that sLNM’s PC1 artifact drives spurious replicability between studies with unrelated disease networks, and that this can be partially mitigated by controlling for PC1 in the permutation test statistic. These simulation results are consistent with our empirical findings. Both the Depression-TMS and Broca’s aphasia sLNM maps showed strong spatial similarity to the PC1 map (Depression-TMS |r| = 0.65; Broca’s aphasia |r| = 0.58) and these two sLNM maps converged significantly even after using the symptom-label permuting null (r = 0.56, p = 0.03). When we used partial correlation controlling for PC1, the two sLNM maps no longer converged (partial r_PC1_ = 0.30, p = 0.11).

### 3.5 Non-specificity of Clinical Outcome Prediction due to PC1

Using a leave-one-subject-out validation scheme (**Fig. 6A**), sLNM predicted TMS response in the Boston TMS cohort (|r| = 0.402) (**Fig. 6B**). However, control maps predicted TMS response equally well or better (**Fig. 6C**). Across 22 control maps, all but two — aphantasia (|r| = 0.262) and political intensity (|r| = 0.187) — predicted outcomes above the level of the cohort’s own leave-one-subject-out sLNM map. The best predictor was the migraine map (|r| = 0.478), an LNM map encoding the connectivity overlap of gray matter atrophy coordinates associated with migraine (Burke et al., 2020; coordinates originally reported by Jia & Yu, 2017). Maps from conditions as diverse as MS-depression (|r| = 0.475), aphasia (|r| = 0.466), the convergent depression map (|r| = 0.462), psychosis (|r| = 0.460), addiction remission (|r| = 0.458), ASD (|r| = 0.457), creativity (|r| = 0.453), OCD (|r| = 0.442), tics (|r| = 0.437), schizophrenia (|r| = 0.431), epilepsy (|r| = 0.430), criminality (|r| = 0.424), the dysphoric-anxiosomatic TMS target (|r| = 0.423), substance use disorder (|r| = 0.422), mania (|r| = 0.420), tremor (|r| = 0.419), PTSD (|r| = 0.412), and Broca’s aphasia (|r| = 0.406) Crucially, the degree map — derived entirely from the healthy GSP1000 connectome with no patient information — also outperformed the cohort’s own sLNM map (|r| = 0.448).

After controlling for PC1, control maps from unrelated conditions that previously outperformed the cohort’s own sLNM map showed near-zero or substantially reduced prediction: tremor (|r| = 0.010), epilepsy (|r| = 0.024), tics (|r| = 0.043), schizophrenia (|r| = 0.049), substance use disorder (|r| = 0.086), Broca’s aphasia (|r| = 0.086), mania (|r| = 0.096), OCD (|r| = 0.098), the degree map (|r| = 0.106), ASD (|r| = 0.131), creativity (|r| = 0.137), and addiction remission (|r| = 0.146). Meanwhile, the strongest predictors are the dysphoric-anxiosomatic TMS target (|r| = 0.264), followed by MS-depression (|r| = 0.259) and the cohort’s own leave-one-subject-out TMS map (|r| = 0.256) — these maps were all derived from cohorts of subjects with depressive symptoms (Siddiqi et al., 2020, 2023) — suggesting that some disease-specific signal related to depression is retained after PC1 is removed.

Controlling for PC2 or PC3 did not improve prediction specificity (Supplementary **Fig. S4**), suggesting that PC1 is the primary driver of the non-specificity in clinical outcome prediction.

## 4 Discussion

LNM and sLNM both offer practical approaches for studying the network effects of focal brain abnormalities. sLNM in particular has been proposed as a method for causal network mapping, because when the same network predicts symptom severity from lesion data and symptom improvement from stimulation data, the convergence of these opposing directional relationships supports causal inference (Siddiqi et al., 2022). We began this investigation to validate sLNM as a more robust extension of LNM. Using simulated datasets with known lesion–symptom relationships and ground truth networks, we expected sLNM to closely approximate ground truth (Petersen et al., 2026; Siddiqi et al., 2026; Zalesky & Cash, 2026). Instead, our findings, together with those of van den Heuvel and colleagues, suggest that sLNM maps may contain systematic bias from the normative connectome. However, as we discuss below, these methodological limitations may partly be addressed with appropriate null models and improved statistical implementation.

To understand how the method can be improved, it is important to differentiate conventional LNM (LNM hereafter) from sLNM. Van den Heuvel and colleagues found that LNM converged towards degree, because LNM identifies overlapping regions functionally connected to the same set of lesions, preferentially including a restricted set of hub regions with widespread connections. However, as argued by the creators of LNM, these maps represent only the sensitivity test—the first step in LNM (Siddiqi et al., 2026). The subsequent specificity test, which contrasts disorder lesions against control lesions, has been demonstrated to increase specificity in LNM findings (for further discussion on LNM specificity testing, see Petersen et al., 2026 and Siddiqi et al., 2026)

In contrast, sLNM does not employ the same specificity test contrasting different disorder groups.In contrast, sLNM does not employ the same specificity test contrasting different disorder groups. Where an LNM study of depression would contrast lesions causing depression against control lesions (Padmanabhan et al., 2019), sLNM can instead correlate symptom scores with connectivity across a clinically heterogeneous lesion dataset (such as the Vietnam Head Injury dataset; Raymont et al., 2011), ranging from no depression to severe depression (Siddiqi et al., 2021). Here, each voxel’s correlation value with depression score replaces the need to contrast different groups. Specificity is further established through cross-dataset replication, showing that independent datasets measuring the same symptom converge onto the same map after symptom-label permutation testing. Unfortunately, our findings showed that sLNM produced maps with amplified PC1 features compared to ground-truth, and this systematic bias is not accounted for in the current null test. The reason for this convergence is mathematical. The PC1 map is, by definition, the spatial pattern explaining the most variance in FC across brain regions. In a typical sLNM analysis, lesions or stimulation targets vary anatomically and therefore span a range of positions along this axis. When the connectivity profiles of these lesions are correlated with symptom scores, the resulting map will inevitably contain a PC1 component, because PC1 captures the largest source of variance along which these connectivity profiles differ (for further discussion, see Supplementary S9 of van den Heuvel et al.). Breaking the structured lesion–symptom pairing, as done in the permutation null, removes the systematic bias towards PC1 in the null distribution, and therefore does not account for PC1-driven convergence between datasets.

For example, two datasets with spatially distinct ground truth networks (r = 0.01) — nonetheless produce sLNM maps that share enhanced PC1 features which drives spatial similarity **(Fig. 5A)**, leading to the incorrect conclusion that both datasets converged onto the same disease network. The problem is further compounded by the common step averaging the sLNM maps together across replication datasets in an attempt to uncover common features between the two maps, which risks further emphasizing PC1 instead of disease-specific features (e.g., “Convergent Broca-Depression Network” in **Fig. 1C**; |r| with PC1 = 0.69). Under a PC1-controlled null, however, this similarity is no longer significant. We confirmed this empirically: maps for TMS depression response and Broca’s aphasia no longer converged after controlling for PC1. Our simulations also showed that controlling for PC1 retains specificity — datasets with spatially similar ground truths still converged — though some sensitivity is lost at lower effect sizes (η² = 0.3). However, this partial correlation approach only partially mitigates the problem, and more robust null models are needed. Regardless, we recommend that future sLNM studies account for PC1 in their null tests.

The next point concerns the clinical utility of these network maps. Most sLNM studies establish the utility of their maps via clinical prediction using leave-one-out validation schemes. When multiple symptoms or clinical outcomes are available, specificity is demonstrated by showing that a given map best predicts the symptom of interest (e.g., depression) versus other symptoms (e.g., sleep, motor function) (Milano et al., 2025; Siddiqi et al., 2023, 2025; Siddiqi, Klingbeil, et al., 2024). Here we tested a different kind of specificity, where the symptom being predicted is the same (TMS response in depression) while varying the predictor map instead. We found that control maps from unrelated disorders, and even the degree map, outperformed the disorder-specific sLNM map. Controlling for PC1 in the validation scheme partially improved specificity, suggesting that a significant proportion of reported clinical prediction may be partly attributable to PC1 features within these maps rather than disease-specific features. We therefore recommend that establishing true specificity requires not only showing that an sLNM map specifically predicts the symptom of interest — a step already performed in sLNM studies — but also that the disease-specific sLNM map outperforms other non-specific maps. Because many of these maps contain PC1 features (or degree features in the case of LNM), apparent clinical prediction may be non-specific, and maps should demonstrate predictive capability even after accounting for PC1.

This leads into an important discussion: why would PC1 have clinical utility in the first place? One possible explanation is that PC1 reflects a biologically meaningful axis in the brain. It closely approximates the principal gradient, the fundamental organizational axis spanning from sensorimotor/unimodal cortex to association/transmodal cortex (Margulies et al., 2016). Lesions at opposite ends of this axis would plausibly affect symptoms differently, given that the sensorimotor and association cortex differ not only in function—from basic sensorimotor processing to abstract cognition (Margulies et al., 2016) — but also in cortical morphology, hubness, cytoarchitecture, gene expression, metabolism, and vulnerability to pathology, role in development of functional brain networks, and responses to brain stimulation (Fornito et al., 2025; Iaccarino et al., 2021; Lowe et al., 2019; Luo et al., 2024; Momi et al., 2025; Vogel et al., 2023). Critically, our findings do not invalidate the LNM framework itself. Studying the remote effects of lesions or treatments in network space, beyond anatomical location alone, remains a central goal of connectomics and network neuroscience (Fornito et al., 2015; Fox, 2018; Vogel et al., 2023). Notably, the bias toward PC1 occurred only when we modeled meaningful lesion–symptom relationships (η² > 0), not when lesions and symptoms were random (η² = 0). Convergence between datasets in the sLNM literature therefore likely reflects genuine network effects within the datasets. The problem is not that the framework is wrong, but that the method inevitably amplifies PC1 features in the output, confounding the disease-specific signal. Importantly, controlling for PC1 in validation steps reduces false positives but does not affect the output map itself. Further work is therefore needed for the method to faithfully reconstruct ground truth. One avenue may be to regress out PC1 features, though this assumes PC1 is not relevant to the disease of interest—for example, regressing out PC1 may falsely remove default mode network nodes if these are part of the true disease network. Another approach may be to modify the correlation step itself, using a general linear model to relate lesion voxel FC and symptoms while accounting for connectome features (PC1-3, and topological degree) as covariates. Finally, this study and previous LNM critiques have emphasized the importance of ground truth simulations for benchmarking (van den Heuvel et al., 2026; Zalesky & Cash, 2026). sLNM is particularly difficult to benchmark because one must simulate both lesions and symptoms together. Here we used a disease model based on unthresholded brain map spatial similarity, as validated in previous studies (Ferguson et al., 2024; Goede et al., 2025; Horn et al., 2017; Li et al., 2021; Siddiqi et al., 2021, 2023, 2025; Sobesky et al., 2022), but future work may incorporate different disease models—for instance, symptoms could be modeled based on connectivity strength to a single region of interest rather than a whole-brain map (Fox et al., 2012), or on the overlap between lesion location and network boundaries (Reich et al., 2022; Siddiqi, Klingbeil, et al., 2024). Testing whether the method can reconstruct latent factors underlying these diverse simulated datasets will be essential for establishing its validity.

In conclusion, sLNM is a promising method grounded in sound theory, but its current implementation systematically overestimates PC1 features which may lead to spurious similarities between disorders. Performing partial correlation to control for PC1 may offer a partial remedy to improve disease-specificity while reducing false positives. Further work is needed to refine sLNM such that the output maps themselves faithfully reconstruct disease-specific networks without bias. The clinical utility of previous sLNM maps may remain intact, but their interpretation would benefit from integration with the growing literature on the sensorimotor–association gradient of cortical organization.

## Acknowledgements

We thank Martijn P. van den Heuvel for conceptual discussions and feedback.

## Author contributions

Each author has made a significant contribution to the manuscript and all authors read and approved its final version. **S.T.**: Conceptualization, Data curation, Formal analysis, Investigation, Methodology, Software, Visualization, Writing — original draft, Writing — review & editing; **A.K.**: Writing — original draft, Writing — review & editing, Software; **S.J.**: Conceptualization, Methodology, Writing — original draft, Writing — review & editing:. **W.P.**: Conceptualization, Methodology, Validation, Software; **A.C.**: Conceptualization, Supervision, Writing — review & editing; **S.S.**: Methodology, Validation; **J.B.**: Supervision, Writing — review & editing; **C.C.**: Conceptualization, Supervision, Funding acquisition, Methodology, Project administration.

## Data availability

All data used in the present study are publicly available. The preprocessed normative fMRI data from the GSP1000 dataset are available at https://doi.org/10.7910/DVN/ILXIKS. Preprocessed lesion files of Broca’s aphasia subjects are available from GitHub (https://github.com/neurolabusc/AphasiaRecoveryCohortDemo). MNI coordinates of TMS sites and their respective improvement in depression are available from the original publication (Weigand et al., 2018). The Yeo-Schaefer atlas was downloaded from GitHub (https://github.com/ThomasYeoLab/CBIG).

## Ethics

This study used publicly available, anonymised datasets. Ethical standards for the original data collection are described in the respective source publications.

## Code availability

Analysis code is provided on GitHub (https://github.com/SasinJamesCCCN/symptom_lnm)

## Use of Generative Artificial Intelligence Assistance

Artificial intelligence (AI) tools were used to assist with minor grammar and phrasing refinements of the text. All AI-suggested edits were reviewed and approved by the authors. All figures were created and all citations were sourced by the authors without AI assistance.

