## Supplementary Material for "Connectome Principal Component Drives Cross-Dataset Replication and Clinical Prediction in Symptom Lesion Network Mapping"

### Figure S1. PC1 amplification of sLNM pipeline

Each scatterplot shows 300 pairs of sLNM maps and their corresponding ground-truth maps, plotting the spatial correlation (Pearson's r) of each map with the first three connectome PCs (top row: PC1; middle row: PC2; bottom row: PC3). Columns represent increasing effect size, from $\eta^2 = 0.0$ (random) on the left to $\eta^2 = 0.99$ (deterministic) on the right. As effect size increases, sLNM–PC1 correlations are amplified relative to their ground-truth–PC1 values, whereas PC2 and PC3 correlations remain closely matched to ground truth across all effect sizes.

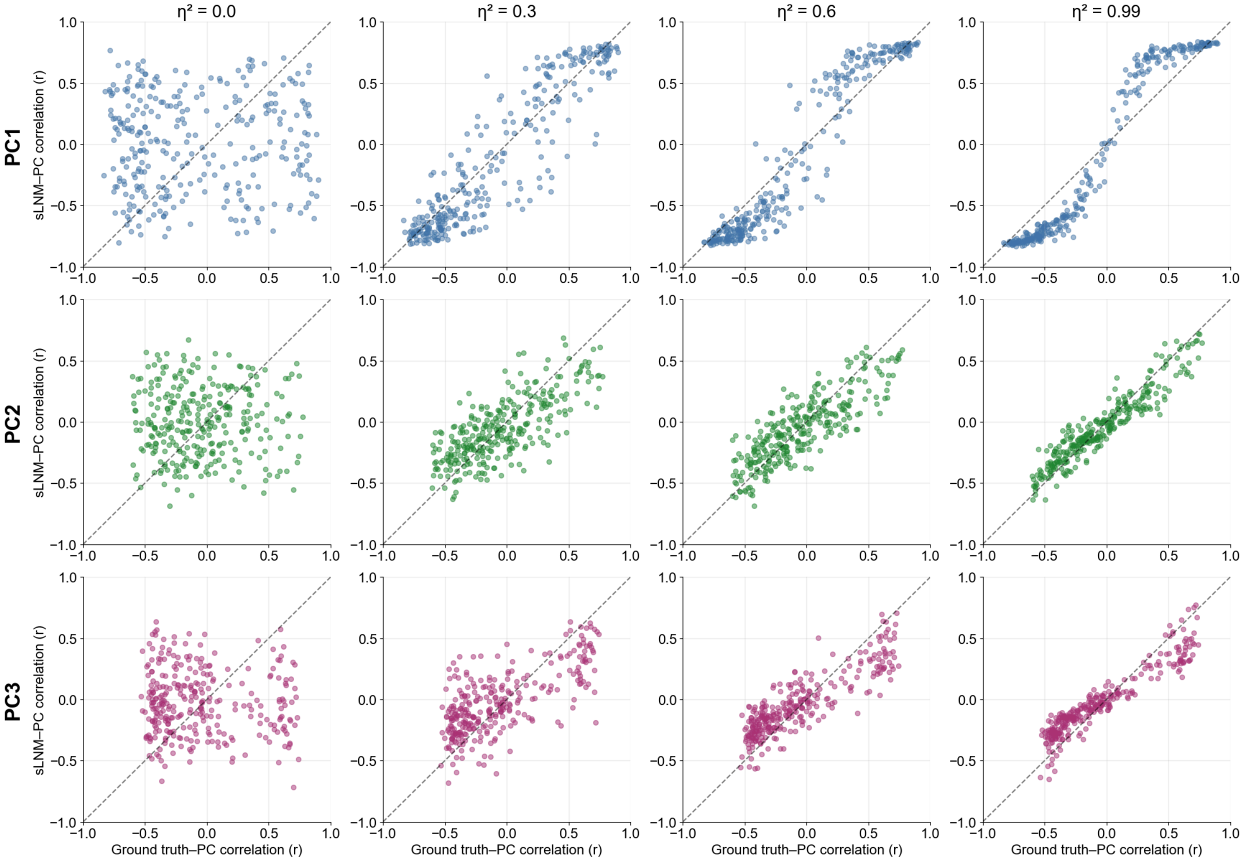

### Figure S2. Controlling for PC2 or PC3 does not affect test statistics

Columns represent effect sizes $\eta^2 = 0.3$ (left), 0.6 (middle), and 0.99 (right). Rows 1–2: using partial correlation controlling for PC2 as the test statistic does not change the significance rate across ground-truth spatial correlation bins (row 1) or the p-value distribution under symptom-label permutation (row 2, QQ plot). Rows 3–4: the same holds when controlling for PC3. Together, this supports the conclusion that PC1, rather than higher-order PCs, drives the spurious convergence between datasets.

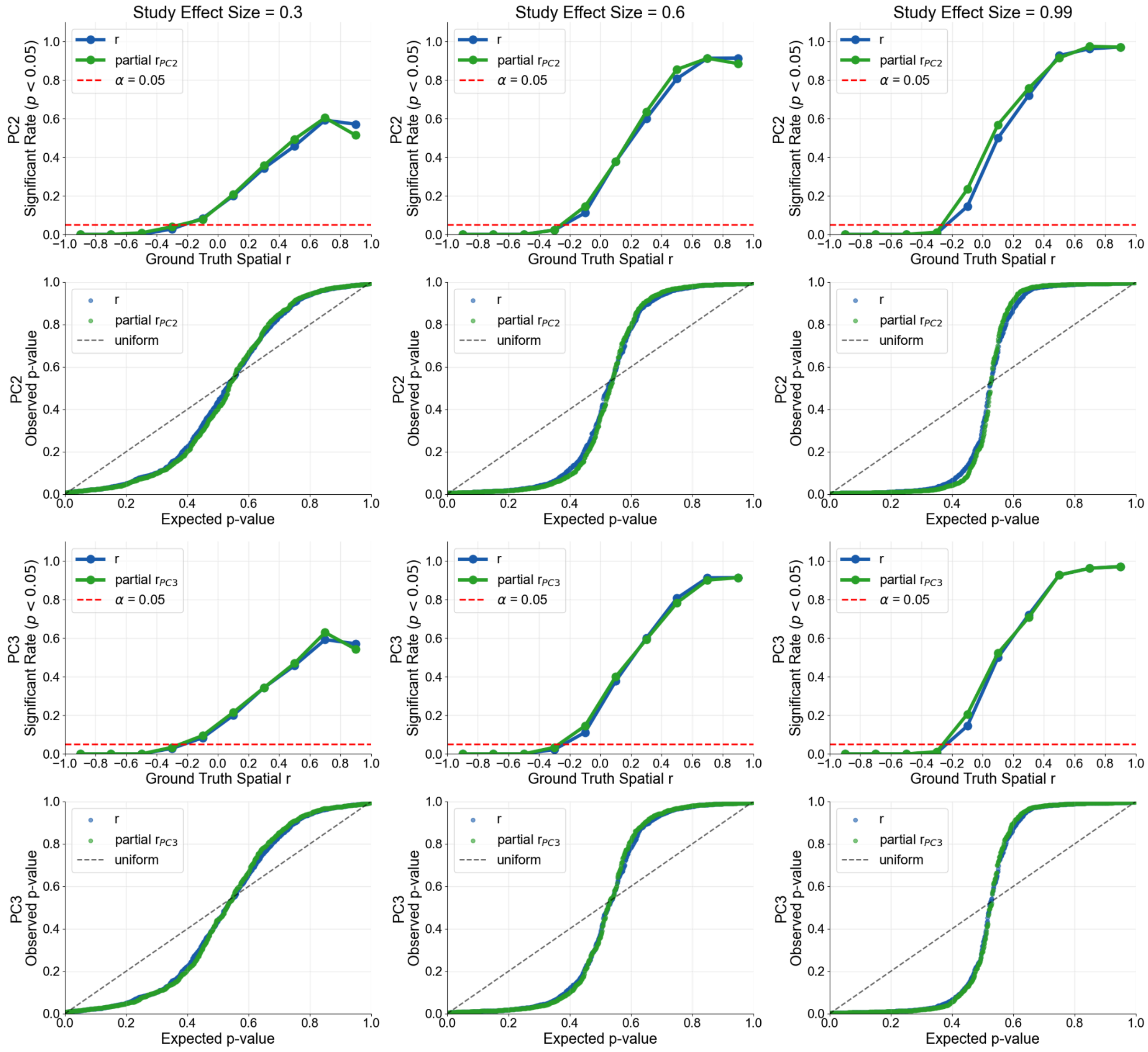

**Figure S3. Visualization Control Maps**

Control networks used for TMS response prediction. Voxel-wise statistical maps for the control networks used to predict TMS response in the Boston TMS cohort (excluding the Broca's aphasia network shown in Fig. 1A of main text). Maps were projected onto the fsLR-32k very-inflated surface for visualization.

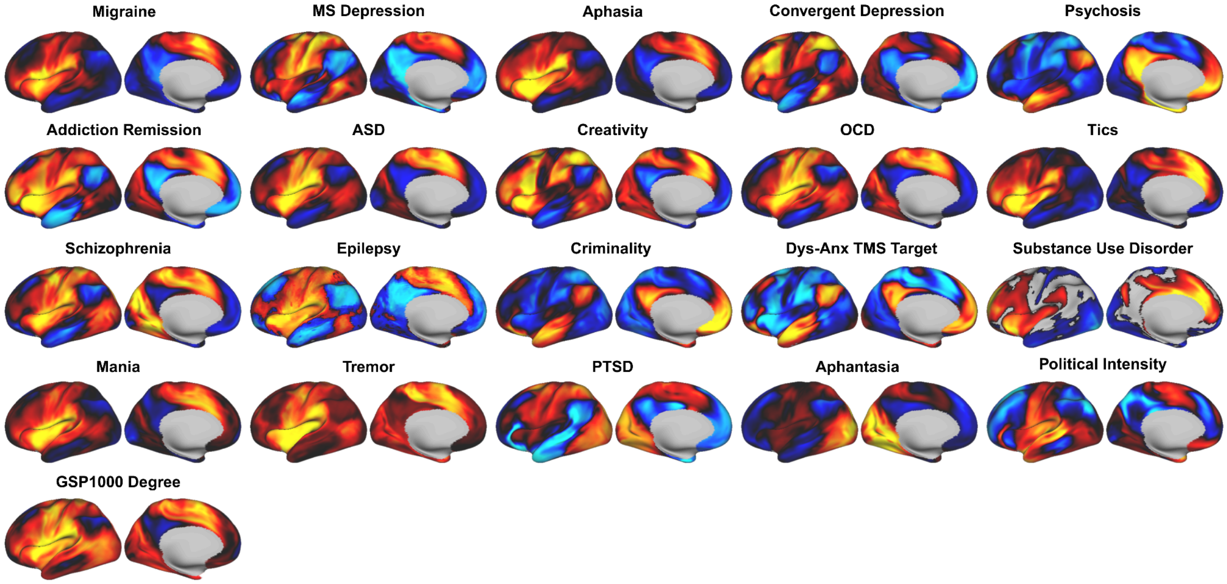

**Figure S4. Prediction of TMS response using spatial similarity to the predictor map versus partial similarity controlling for PC1, PC2, or PC3. Column 1:** prediction using raw spatial similarity to the leave-one-subject-out sLNM cohort map shows all but two control maps outperforming the cohort map. **Column 2:** controlling for PC1 substantially improves specificity — the cohort map becomes the third-best predictor, closely following the Dysphoric-Anxiosomatic TMS target and MS depression maps. **Columns 3 and 4:** controlling for PC2 or PC3, respectively, does not improve specificity, with all but three control maps still outperforming the cohort map.

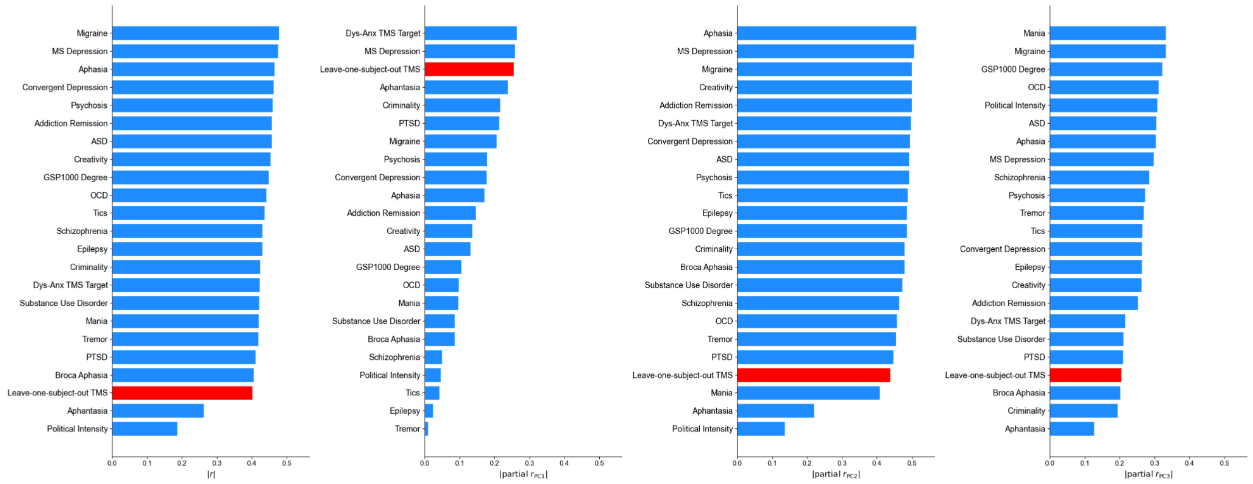

**Supplementary Table S1** Published sLNM maps (N = 5) used as control maps to predict TMS response. For each map, the table lists the map name, the population or input lesions used to derive it, the sLNM method applied, the source of the map, the map quantity (statistical value represented), and the original publication.

| Map Name | Population/Input | Method | Map Source | Map Quantity | Reference |
|---|---|---|---|---|---|
| Dysphoric-Anxiosomatic Target | Depression subjects receiving TMS for treatment of depression (N=111 for active TMS group) | **sLNM:** FC from each TMS seed correlated with individual symptom scores, yielding 21 maps (21 item-BDI scale) and 28 maps (28 item-HAM-D scale).<br>**Clustering:** sLNM maps clustered by spatial correlation into two groups (anxiosomatic, dysphoric); averaged within clusters, then combined into a composite atlas (clusters are near sign-flipped versions of each other). | https://neurovault.org/images/787858/ | Positive values are hypothesized to be better TMS targets for anxiosomatic symptoms. Negative values are hypothesized to be better TMS targets for dysphoric symptoms. | (Siddiqi et al., 2020) |
| Convergent Depression | Brain lesions (N=461, five datasets), TMS treatment for depression (N=151, four datasets), DBS (N = 101, five datasets) | **sLNM**: seed (lesion or TMS or DBS) FC value correlated with depression score in each cohort. Composite map derived from 14 datasets reported. | https://neurovault.org/images/787859/ | Connectivity to all combined modalities (lesion, TMS, DBS) which correlated with depression severity | (Siddiqi et al., 2021) |
| MS Depression | Multiple sclerosis (N=281) | **sLNM**, seed (MS white matter lesion) FC connectivity correlated with Neuro-QoL depression subscale | https://neurovault.org/images/787866/ | Connectivity to MS lesion which correlated with Neuro-QoL depression subscale | (Siddiqi et al., 2023) |
| PTSD | Vietnam Head Injury Study (Raymont et al., 2011) subjects with penetrating head trauma (N=193) | **sLNM:** Lesion FC values correlated with PTSD status | https://neurovault.org/collections/20627/ | Connectivity to lesion which lowers risk of PTSD | (Siddiqi et al., 2024) |
| Political intensity | Vietnam Head Injury Study (Raymont et al., 2011) subjects with penetrating head trauma (N=124) | **sLNM:** Lesion FC values correlated with intensity of political involvement | https://neurovault.org/collections/20530/ | Connectivity to lesion which correlated with greater intensity in political involvement | (Siddiqi et al. 2025) |

**Supplementary Table S2** Published LNM maps and methodological variants (N = 4) used as control maps to predict TMS response. For each map, the table lists the map name, the population or input lesions used to derive it, the LNM method applied, the source of the map, the map statistic, and the original publication.

| Map Name | Population/Input | Method | Map Source | Map Quantity | Reference |
|---|---|---|---|---|---|
| Tremor | Lesions relieving tremors (N=11) | **LNM**: Lesion as seeds | Lead-DBS - Essential Tremor Lesion Network Atlas (Joutsa 2018) | Overlap of FC values of all lesions relieving tremors | (Joutsa et al., 2018) |
| Substance Use Disorder | MNI coordinates of brain atrophy (N=45 studies) and fMRI abnormalities (N=99 studies) | **Coordinate Network Mapping:** MNI coordinate as seeds | https://github.com/nimlab/NMH_Stubbs2023/blob/master/SUD_CNM_FINAL.nii.gz | Overlap of FC values of all MNI coordinates for atrophy and fMRI abnormalities | (Stubbs et al., 2023) |
| Epilepsy | Idiopathic generalized epilepsy (IGE): **1)** 131 MNI coordinates from 21 studies (brain atrophy or fMRI hyperactivity studies) and **2)** 21 subjects with IGE receiving centromedial nucleus (CM) DBS | 1) **Coordinate Network Mapping:** LNM performed on MNI coordinates (IGE map)<br>2) **modified LNM*:** FC map of DBS seeds average into a single map, weighted by each subject % reduction in seizure frequency (CM DBS) | https://github.com/jigongjun/IGENetwork/blob/main/IGEnetwork.nii | Composite z-score map from averaging 1) IGE map and 2) CM DBS map together | (Ji et al., 2025) |
| Psychosis | Brain lesions causing psychosis (N=153) | **LNM:** brain lesion as seed | https://neurovault.org/images/899080/ | Overlap of FC value of all psychosis lesions | (Pines et al., 2025) |

**Supplementary Table S3** LNM maps (N=11) and the GSP1000 degree map used as control maps to predict TMS response. These LNM maps were computed by van den Heuvel and colleagues and rather than the original LNM authors themselves, using seed coordinates, cortical deviations, or lesions reported in the original publications.

| Map Name | Population/Input | Method | Map Source | Map Quantity | Reference |
|---|---|---|---|---|---|
| Criminality | Lesions causing criminal behavior (N=17) | **LNM:** Lesion as seeds | https://github.com/dutchconnectomelab/lesionnetworkmapping/blob/main/data/LNM_networks/computed_LNM_lesions/DARBY_CRIMINALITY_2mm.nii.gz | Overlap of FC value of all lesions causing criminality | (Darby et al., 2018) |
| Migraine | Meta-analysis of migraine voxel-based morphometry studies (Jia and Yu, 2017) | **Coordinate network mapping** (**LNM** performed on reported 11 MNI coordinates associated with cortical deviations in migraine studies) | https://github.com/dutchconnectomelab/lesionnetworkmapping/blob/main/data/LNM_networks/computed_LNM_coordinates/BURKE_MIGRAINE_2mm.nii.gz | Overlap of FC value of all migraine MNI coordinates | (Burke et al., 2020) |
| Mania | Lesions causing mania (N=41) | **LNM**: Lesion as seeds | https://github.com/dutchconnectomelab/lesionnetworkmapping/blob/main/data/LNM_networks/computed_LNM_prevalence/COTOVIO_MANIASETA_2mm.nii.gz | Overlap of FC value of all lesions causing criminality | (Cotovio et al., 2020) |
| Tics | Lesions causing Tics (N=22) | **LNM:** Lesion as seeds | https://github.com/dutchconnectomelab/lesionnetworkmapping/blob/main/data/LNM_networks/computed_LNM_lesions/GANOS_TICS_2mm.nii.gz | Overlap of FC value of all Tic lesions | (Ganos et al., 2022) |
| Addiction Remission | Brain lesion in smokers resulting in remission or no remission of smoking behavior (N=129) | **Group contrast LNM:** general linear model (GLM) performed on lesion FC value and subject group label (remission vs no-remission) | https://github.com/dutchconnectomelab/lesionnetworkmapping/blob/main/data/LNM_networks/computed_LNM_lesions/JOUTSA_ADDICTIONCONTRASTAB_2mm.nii.gz | GLM weights of FC value predicting remission in smoking behavior | (Joutsa et al., 2022) |
| ASD | Extreme cortical deviations in Autism Spectrum Disorder (N=202) | **LNM**: Individual cortical deviations as seeds | https://github.com/dutchconnectomelab/lesionnetworkmapping/blob/main/data/LNM_networks/computed_LNM_cort_dev/SEGAL_ASD_2mm.nii.gz | Overlap of FC value of all ASD cortical deviations | (Segal et al., 2023) |
| OCD | Extreme cortical deviations in OCD (N=167) | **LNM**: Individual cortical deviations as seeds | http://github.com/dutchconnectomelab/lesionnetworkmapping/blob/main/data/LNM_networks/computed_LNM_cort_dev/SEGAL_OCD_2mm.nii.gz | Overlap of FC value of all OCD cortical deviations | (Segal et al., 2023) |
| Schizophrenia | Extreme cortical deviations in schizophrenia (N=383) | **LNM**: Individual cortical deviations as seeds | https://github.com/dutchconnectomelab/lesionnetworkmapping/blob/main/data/LNM_networks/computed_LNM_cort_dev/SEGAL_SCZ_2mm.nii.gz | Overlap of FC value of all schizophrenia cortical deviations | (Segal et al., 2023) |
| Aphasia | Post-Stroke Aphasia (N=20) *** used as control lesions, main study focused on apraxia population)* | **LNM:** stroke lesion as seed | https://github.com/dutchconnectomelab/lesionnetworkmapping/blob/main/data/LNM_networks/computed_LNM_coordinates/KUTCHE_CREATIVITY_2mm.nii.gz | Overlap of FC value of all aphasia lesions | (Zarifkar et al., 2023) |
| Creativity | Meta-analysis of brain coordinates activated by creativity in task fMRI (36 studies) | **Coordinate network mapping**: LNM performed on 36 study coordinate seeds | https://github.com/dutchconnectomelab/lesionnetworkmapping/blob/main/data/LNM_networks/computed_LNM_coordinates/KUTCHE_CREATIVITY_2mm.nii.gz | Overlap of FC value of all MNI coordinates activated in creativity task fMRI studies | (Kutsche et al., 2025) |
| Aphantasia | Lesions causing aphantasia (N=12) | **LNM:** Lesion as seeds | https://github.com/dutchconnectomelab/lesionnetworkmapping/blob/main/data/LNM_networks/computed_LNM_lesions/KUTSCHE_APHANTASIA_2mm.nii.gz | Overlap of FC values of all lesions relieving tremors | (Kutsche et al., 2026) |
| GSP1000 Degree | Healthy GSP1000 connectome subjects (N=1000) | Compute degree (row sum of group-averaged connectivity matrix) | https://github.com/dutchconnectomelab/lesionnetworkmapping/blob/main/data/degree/GSP1000_degree.nii.gz | Average functional degree of normative connectome | (van den Heuvel et al., 2026) |
